# PEP-Patch: Electrostatics in Protein-Protein Recognition, Specificity and Antibody Developability

**DOI:** 10.1101/2023.07.14.547811

**Authors:** Franz Waibl, Nancy D. Pomarici, Valentin J. Hoerschinger, Johannes R. Loeffler, Charlotte M. Deane, Guy Georges, Hubert Kettenberger, Monica L. Fernández-Quintero, Klaus R. Liedl

**Affiliations:** Department of General, Inorganic and Theoretical Chemistry, and Center for Molecular Biosciences Innsbruck (CMBI), University of Innsbruck, Innsbruck, Austria; Department of Statistics, University of Oxford, Oxford, UK; Roche Pharma Research and Early Development, Large Molecule Research, Roche Innovation Center Munich, Penzberg, Germany

**Keywords:** Electrostatic potential, molecular recognition, electrostatic patches, substrate specificity, antibody developability, affinity maturation

## Abstract

The electrostatic properties of proteins arise from the number and distribution of polar and charged residues. Due to their long-ranged nature, electrostatic interactions in proteins play a critical role in numerous processes, such as molecular recognition, protein solubility, viscosity, and antibody developability. Thus, characterizing and quantifying electrostatic properties of a protein is a pre-requisite for understanding these processes. Here, we present PEP-Patch, a tool to visualize and quantify the electrostatic potential on the protein surface and showcase its applicability to elucidate protease substrate specificity, antibody-antigen recognition and predict heparin column retention times of antibodies as an indicator of pharmacokinetics.

## INTRODUCTION

Every protein consists of a unique combination of amino acids, which exhibit distinct biophysical properties and therefore determine the life cycle of a protein, from folding to biological function and degradation.^1–4^

Electrostatic forces are central in molecular biology, co-determining protein folding, protein-protein interactions, protein-DNA/RNA interactions, ion binding, dimerization, and protein stability.^1,2,4^ Additionally, they affect the pKa values of ionizable groups in proteins and DNA/RNA.^5,6^ The complex multi-step process of molecular recognition requires a balance of entropic and enthalpic components.^7,8^ Enthalpic and entropic contributions to binding vary between the different steps from unbound to the fully bound state. Electrostatics are a critical component of the enthalpic contributions, dominating and guiding the recognition process, especially when the binding partners are still distant from one another.^9^ Proteases in particular have been shown to have an electrostatics-driven recognition process, as the substrate preferences can be predicted from charge complementarity in the binding interface.^9,10^

Electrostatic forces affect molecular binding not only through interactions between the binding partners, but also through interactions with the solvent. This is because solvent molecules must be displaced from the binding interface, which introduces a large desolvation penalty that needs to be overcome by an interplay of attractive electrostatic and hydrophobic interactions upon protein-protein or protein-ligand association. Additionally, it has been reported that higher electrostatics due to their long-range interaction network can increase specificity and at the same time restrict flexibility.^11^ On the other hand, weak electrostatics can be associated with conformational variability and, consequently, cross-reactivity.^11^ Protein folding and thermal stability are also influenced by electrostatics. In particular, polar interactions are a major contributor to hydrogen bonding, and hydration of charged and polar amino acids has a profound impact on correct protein folding.^1,2,4^ Due to protonation state changes, pH can influence protein stability and function.^6,12^ Thus, understanding the role of electrostatics in protein function is crucial to advance, guide and facilitate protein engineering and design.

The high therapeutic potential of antibodies in combination with their versatility makes them excellent candidates to study.^13,14^ Biophysical properties of antibodies such as surface charges or hydrophobicity and the isoelectric point are thought to be responsible for changes in pharmacokinetics, efficacy, dose intervals and application route.^15,16^ One common measure of an important biophysical property is Heparin retention chromatography. Heparin retention chromatography for antibodies has been shown to separate antibodies with different serum half-life.^17^ Heparin is a negatively charged polysaccharide, that resembles the glycocalyx, a saccharide layer on the inside of epithelial cells. Interaction with the glycocalyx is believed to increase the propensity of a compound to be taken up into the cell by pinocytosis, followed by digestion and thus the retention time in heparin chromatography correlates with serum-half-life of monoclonal antibodies.^17^

### Surface patches

Macromolecular interactions are often mediated through a single dominant interaction surface. In the case of primarily electrostatic interactions, this implies that continuous surface patches with a high charge density are likely candidates for interaction surfaces. Similar arguments hold for hydrophobic interactions, which might also be mediated by a single hydrophobic patch.^18^

The electrostatic potential around a protein in solution is routinely calculated using Poisson-Boltzmann or Generalized Born calculations.^19–21^ For visualization of the resulting potential, iso-surfaces can be displayed in standard molecular visualization packages such as PyMOL^22^ or VMD.^23^ However, this visualization is not optimal for quantification of the results, since the potential in the first hydration shell, which is an important indicator of interaction strength, is not visible.

On the other hand, when developing quantitative scores of electrostatics of hydrophobicity surface patches have often been used. A patch is usually defined as a continuous surface that fulfills a certain criterion. For instance, a positive surface patch might be defined as having an electrostatic potential exceeding a given threshold value. Software to search for continuous patches has been developed since the 90s^24,25^ and is implemented in molecular visualization software, e.g., such as MOE^26^. However, those implementations are usually tied to a single use case and not broadly applicable.

Here, we present the Python tool PEP-Patch, which allows a user to search for continuous surface patches. While our tool uses an electrostatic potential from APBS by default, it can be used with any combination of a surface representation and a 3-dimensional potential, thus providing a versatile building block for biomolecular analysis.

## METHODS

### Antibody structure models

We used heparin data from Kraft et al.^17^ for a set of 137 antibodies described in the dataset by Jain et al.^27^

To compare the influence of different antibody conformations on the respective electrostatic potential, we used three antibody structure prediction tools, namely DeepAb^28^, ImmuneBuilder^29^ and MOE^26^, using the default settings, to predict Fv structures. In addition, for 49 of these 137 antibodies^27^ crystal structures were available.

### Electrostatic surfaces

Our PEP-Patch tool (v1) is freely available on Github (https://github.com/liedllab/surface_analyses) under an MIT license.

Our tool uses the Advanced Poisson-Boltzmann Solver (APBS) software to compute the electrostatic potential using the Poisson-Boltzmann equation^19,30,31^. A NaCl-concentration of 0.1 M is used by default, without titrating residues. Smooth molecular surfaces can be defined using a Gaussian surface, a solvent-accessible surface, or a Conolly-type surface. The electrostatic potential is evaluated at every surface vertex using linear interpolation between the neighboring grid voxels. Positive and negative surface patches are defined using an isolevel, by searching for connected components in the graph defined by the triangulated surface. Similar procedures are commonly used to find protein surface patches.

The output of our tool includes the surface and interpolated values in the numpy storage format (npz), as well as a color-coded surface in ply format for visualization in molecular viewers such as PyMol or VMD.^32^

### Quantitative scores for electrostatics

We define five different quantitative scores for the electrostatic properties of a molecule. To do so, we start from the electrostatic potential map obtained from a Poisson-Boltzmann calculation. We then select grid voxels that are solvent accessible, but within a defined distance cutoff from the protein. By default, this distance cutoff is defined to be 10 Å of the protein surface.

The *total* score is defined simply as the integral of the electrostatic potential over that region. In a very simplified view, it can be thought of as the interaction strength with a positively charged substance, given that this substance is evenly distributed in the selected volume.

We also define positive and negative scores, which only include contributions of the positive and negative regions in the electrostatic potential. Again, they can be roughly understood as an interaction strength with a charged substance, this time imagining that the substance is distributed only in the respective part of the volume.

Finally, we define *high* and *low* scores, which are defined in the same way as the *positive* and *negative* ones, except that they include only regions above and below a given electrostatic potential cutoff. Additionally, the tool provides the residues that contribute the most area to an electrostatic patch.

### APPLICATION AND ILLUSTRATIVE EXAMPLES

In this study, we present a tool to quantify/characterize electrostatic surface properties of proteins to elucidate determinants of molecular recognition and identify descriptors that govern pharmacokinetics. Here, we apply our tool to three different case studies, elucidating substrate specificity of proteases based on their surface properties, characterizing molecular recognition upon antibody affinity maturation, and predicting biophysical properties to distinguish and optimize pharmacokinetics of antibodies.

### Protease substrate recognition

Proteases are enzymes that proteolytically cleave peptide bonds and play a key role in a variety of different physiological processes, ranging from complex signaling cascades, blood coagulation, and food digestion, to key aspects of the immune system such as programmed cell death and digestions of cells.^33,34^ These very distinct and broad biological functions require vast differences in specificity and promiscuity.^10,35,36^ While some proteases reveal a high specificity for substrate sequences, others are more promiscuous, cleaving a variety of different substrates. An example would be digestive proteases that cleave food proteins, and thus need to function on many different substrates. Substrate specificity of proteases is conveyed by molecular interactions occurring at the protein-protein interface (protease and substrate) in the binding cleft of the protease.^10^ Here, we use our tool to compare proteins based on their electrostatics and demonstrate its applicability on proteases.

Figure 1A shows the comparison of three proteases differing in their substrate preferences. We calculated the electrostatic potential (Figure 1B) based on the available X-ray structures for all three proteases (PDB accession codes: 1PQ7 for Trypsin, 4CHA for Chymotrypsin and 1FQ3 for Granzyme B) and show that by considering the electrostatic potential around each protease (red represents negatively charged patches, blue positively charged patches), the substrate preference^37^ can be inferred, i.e., Granzyme B shows a preference for negatively charged substrates, Trypsin prefers more positively charged substrates and Chymotrypsin rather uncharged substrates. The positive and negative protein surface patches are illustrated in Figure 1C and the residues contributing most to the absolute value ascribed to an electrostatic patch are provided in SI Table 1.

**Figure 1.**
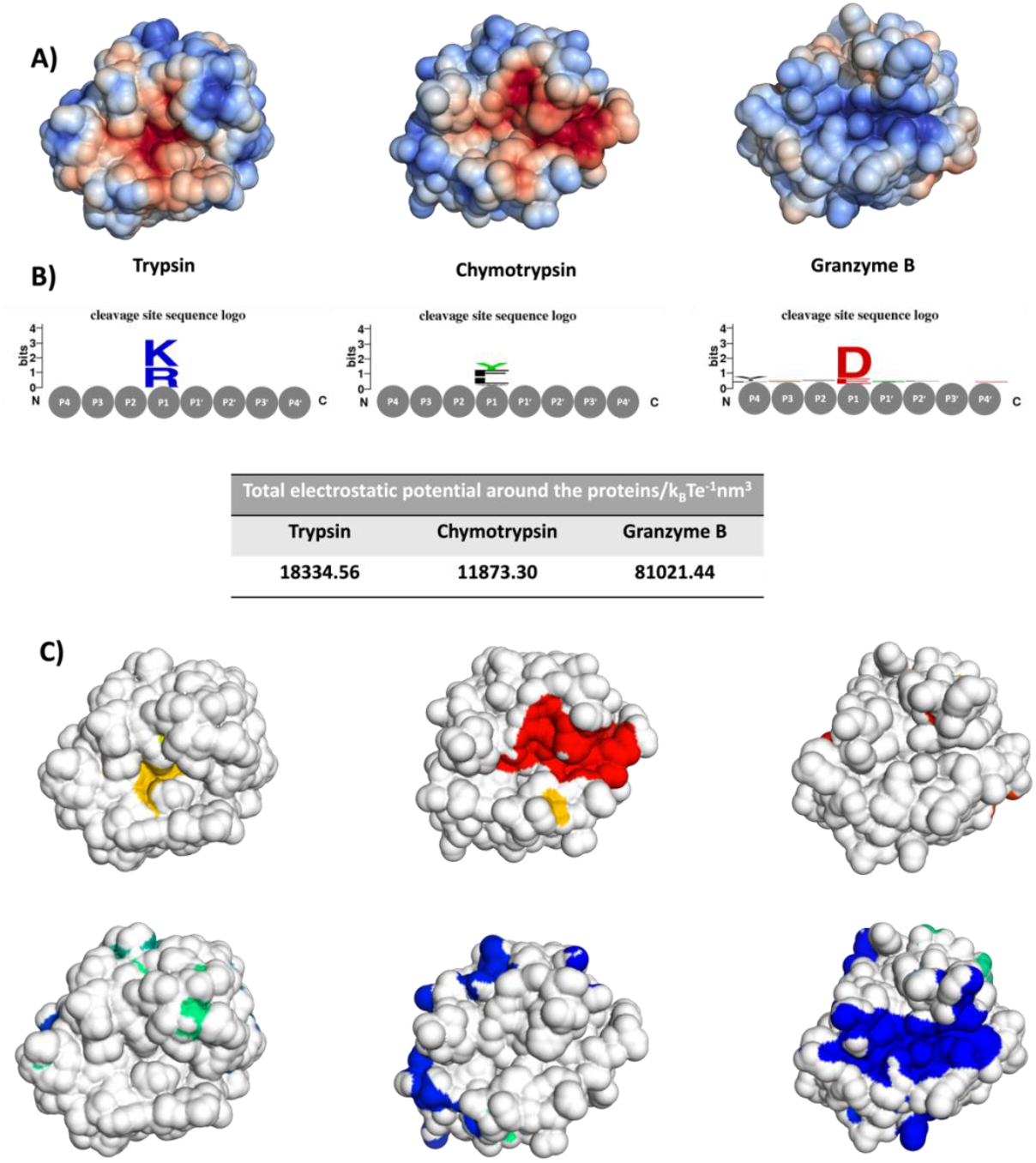
Electrostatic potential around three proteases differing in their substrate preferences. A) The electrostatic potential on the protein surface shows that Trypsin favors positively charged substrates, Chymotrypsin favors rather uncharged substrates and Granzyme B recognizes negatively charged substrates in line with the cleavage site sequences logos obtained from the MEROPS database. B) Summary of the total electrostatic potential around the proteins. C) Positive (blue to green, from biggest to smallest patch) and negative (red to yellow, from biggest to smallest patch) protein surface patches.

### Antibody-antigen recognition

Biomolecular recognition between proteins follows complex mechanisms. Understanding protein-protein interfaces and their interactions is crucial to advance the development of biotherapeutics. Here, we focus on the interface of two different antibodies binding to the same chemokine CXCL13 antigen.^38^ Structurally, the antigen-binding fragment of an antibody (Fab) is composed of a heavy and light chain, which form the antigen-binding site, the paratope. The paratope is primarily found within six hypervariable loops, the complementarity determining region (CDR) loops.^13^ In addition to the CDR loops, residues in the framework as well as the relative interdomain orientation (between the heavy and light chain) can strongly influence antigen recognition.^39–42^ It is well established that for antibodies single-point mutations can result in changes in the binding site conformations and thereby affect biophysical properties.^43–45^ The antibody variants investigated here, the parental 3B4 and the optimized E10, have substantial differences in affinity and stability resulting from four point-mutations located in the CDR-L3 loop.^38^ To calculate the electrostatic potential, we used the available crystal structures (PDB accession codes: 5CBA for 3B4 and 5CBE for E10) and find that these four-point mutations contribute to an improved electrostatic complementarity of the E10 variant with CXCL13. This is reflected in the results presented in Figure 2.

**Figure 2.**
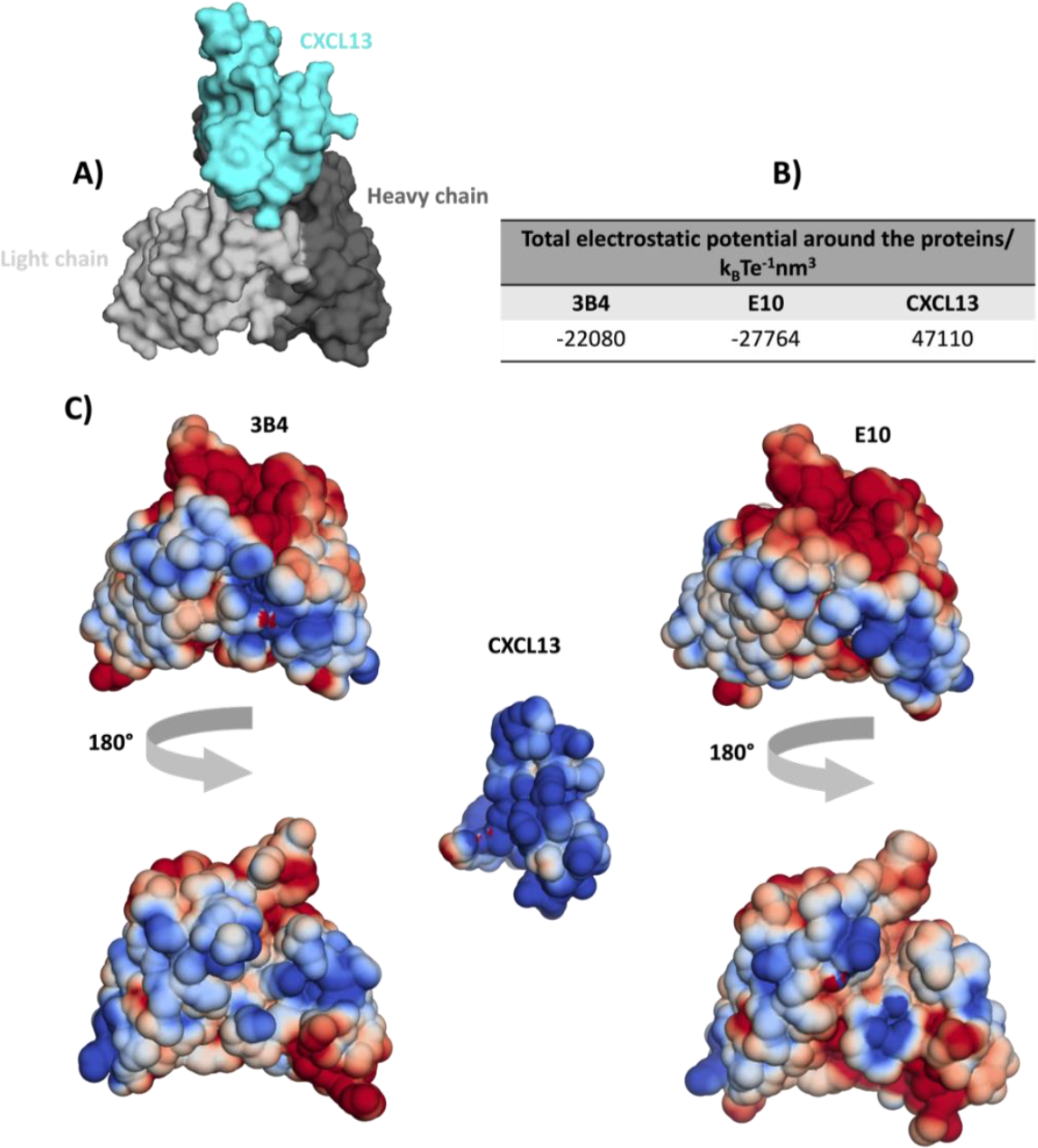
Electrostatic potential in antibody recognition. A) Antibody-antigen binding mode, derived from the available X-ray structure (PDB accession code: 5CBE), showing the antibody in grey and the antigen in cyan. B) Summary of the total electrostatic potential around the proteins. C) Electrostatic potential around the antibodies 3B4 and E10 (both sides) and the CXCL13 antigen.

### Antibody developability – predicting differences in pharmacokinetics

Another critical aspect in developing antibodies apart from antibody-antigen recognition is understanding factors that are responsible for changes in pharmacokinetics. One experimental technique for examining pharmacokinetics are heparin retention chromatography times.^17^

Here, we compared our results to experimentally available relative heparin column retention times and show that the electrostatic potential calculated for the different antibody variable fragments (Fv) is a key determinant for changes in pharmacokinetics, reflected in a compelling correlation with the experiment. We chose to compare three antibody structure prediction tools to understand the conformation dependence of the electrostatic patch prediction. Independent of the structure models or X-ray structures used as starting points of our calculations, we find similar correlations. This is a strong indication that the electrostatic potential, due to its long-ranged nature, may be less conformation dependent than other biophysical properties such as hydrophobicity.

At a low positive electrostatic potential score (roughly below 20 k_B_Te^-1^nm^-1^) there appears to be no correlation with the heparin retention time. This is probably due to very weak interactions with the column. The highlighted points in Figure 3A represent the antibodies with the highest and lowest heparin column retention times. These differences in the experimental retention times are also reflected by the electrostatic potential around the antibodies, i.e., lenzilumab a higher positive electrostatic potential, compared to sirukumab, which is substantially more negative (Figure 3B).

**Figure 3.**
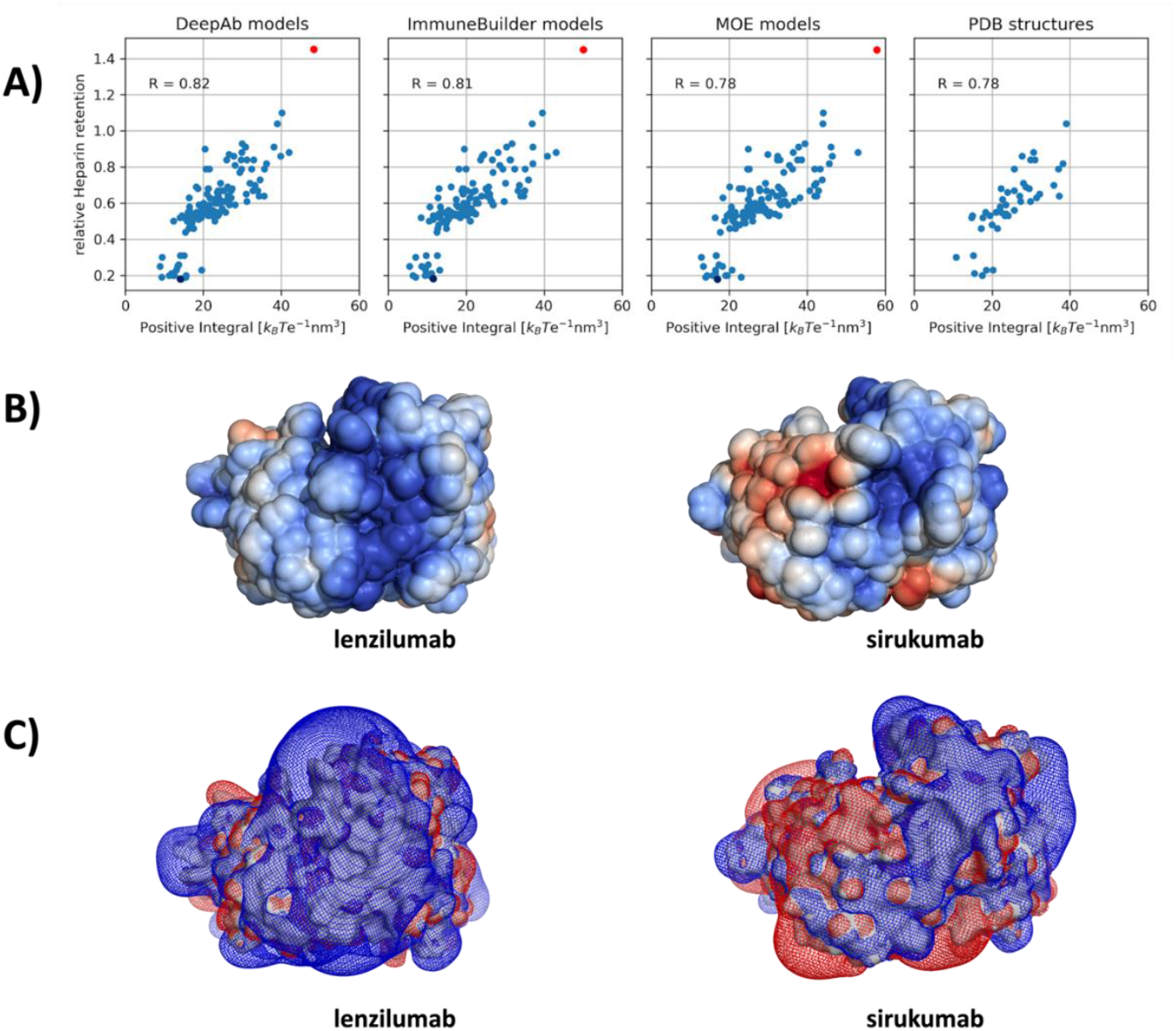
A) Scatter plot of relative Heparin retention time correlated with the integral of the positive electrostatic potential over the solvent-accessible volume, for the investigated 137 antibodies using the DeepAb models, ImmuneBuilder models, MOE models and the available 49 PDB structures. B) Electrostatic potential around the DeepAb models of lenzilumab (depicted as red dot) and the sirukumab (shown as blue dot), which are the antibodies with the highest and lowest Heparin retention time, respectively. C) Mesh isopotential surface around the two DeepAb models of lenzilumab and the sirukumab.

To demonstrate the usefulness of our results, we compare our tool to other commonly used scores for protein charges. We compute the average net charge of each model using the Protein Properties tool in MOE, with the conformational sampling option turned on, and plot the resulting values against the same heparin data (Figure S1). Furthermore, we produce an analogous plot using the total area of positive patches, calculated using the same procedure in MOE (Figure S2). While all three tools perform well in general, we note that the correlation between our tool and the heparin retention times are slightly higher, while the computation is drastically faster, since only a single structure is used for our scores. To make it easier for users to match patch data to protein residues, PEP-patch tool provides the residues that contribute the most area to each patch. This makes it easier to assign patches to protein residues, even without looking at the three-dimensional representation. Furthermore, if the input structure is an antibody fragment, it can detect which patches contain atoms of the complementarity determining regions (CDRs) using ANARCI.^46^

### Other use-cases

To demonstrate the wide applicability of our tool to calculate patches for any user-provided density map, we also show results based on the localized hydration free energy from a grid inhomogeneous solvation theory (GIST)^47–49^ calculation in Figure 4, first presented in Waibl et.al.^50^ Using a solvent-accessible surface at a probe radius of 1.4 Å to approximate the shape of the first hydration shell, we calculate patches for the free energy of hydration around the paratope of bevacizumab (PDB code 1BJ1). We find a larger number of patches showing negative free energy in Figure 4B when compared to the positive patches in Figure 4C, which are rather small in number and size. This is in line with our previous findings that hydrophobicity and IgG antibodies are mainly being found in the serum. For rugged densities such as the free energy on hydration shown here, calculating patches may result in a large number of small-sized patches. For visualization, it is advisable to choose a continuous color map in such cases as the standard qualitative colormap does not provide enough colors to show all patches.

**Figure 4.**
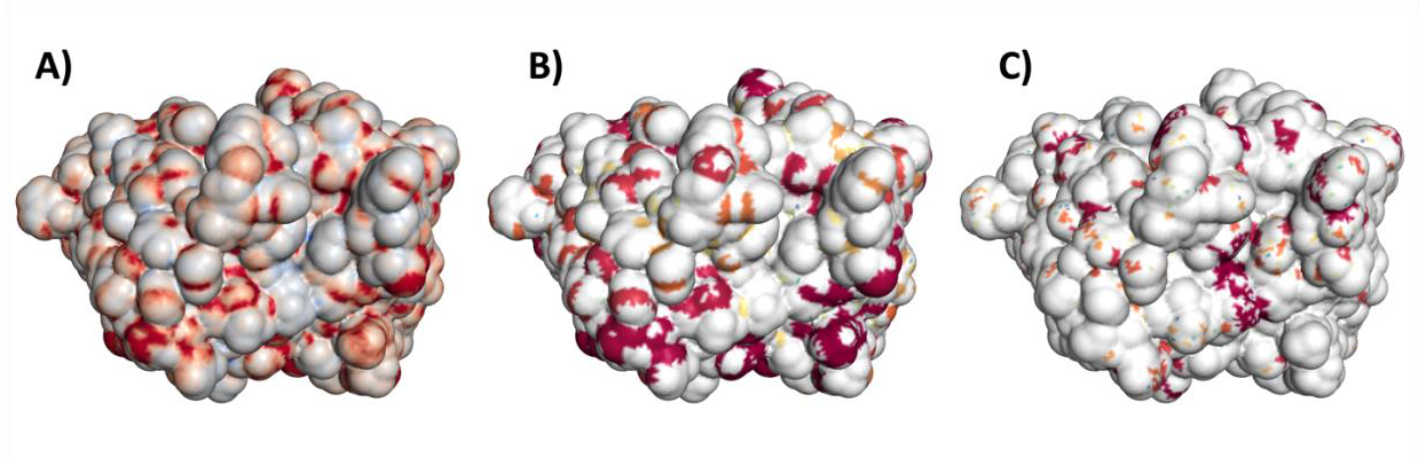
A) Surface of bevacizumab colored by GIST free energy of hydration around the Fv ranging from negative values in red to positive values in blue. B) Patches based on surface vertices with negative free energy of solvation. C) Patches based on surface vertices with positive free energy of solvation. 4 B-C) Patches are colored based on their patch size (from large to small patches colored in red-yellow-green-blue).

## CONCLUSION

We present the PEP-Patch tool, which allows calculating the electrostatic potential of proteins. Here, we show that the electrostatic potential can explain biomolecular recognition, substrate specificity and even pharmacokinetics of antibodies. Additionally, it allows to directly visualize and quantify the electrostatic potential around different proteins, that can guide the design of biotherapeutic proteins. Furthermore, the tool identifies the residues that contribute the most to an electrostatic patch, which can inform rational protein design.

## AUTHOR INFORMATION

The manuscript was written through contributions of all authors. All authors have given approval to the final version of the manuscript. ‡ F.W., N.D.P. and V.J.H. contributed equally.

## FUNDING AND ACKNOWLEDGMENTS

N. D. P. has received funding from the European Union’s Horizon 2020 research and innovation programme under the Marie Skłodowska-Curie grant agreement number 847476. The views and opinions expressed herein do not necessarily reflect those of the European Commission. This work was supported by the Austrian Science Fund (FWF) via the grant P34518. This work was supported by the Austrian Academy of sciences APART-MINT postdoctoral fellowship to M.L.F.Q. We acknowledge CHRONOS for awarding us to access to Piz Daint at CSCS, Switzerland. We acknowledge EuroHPC Joint Undertaking for awarding us access to Karolina at IT4Innovations, Czech Republic.

## ABBREVIATIONS

PB: Poisson-Boltzmann;
Fab: antigen-binding fragment;
CDR: complementarity determining region.

## DATA AND SOFTWARE AVAILABILITY STATEMENT

The code for the PEP-Patch tool is available on Github under the MIT license https://github.com/liedllab/surface_analyses. The structures used in this manuscript are publicly available, with the PDB codes: 1PQ7, 4CHA,1FQ3, 5CBE, 5CBA and the models are available on Github.

## REFERENCES

(1) Sheinerman, F. B.; Norel, R.; Honig, Burrent Opinion in Structural Biology 2000, 10 (2), 153–159. https://doi.org/10.1016/S0959-440X(00)00065-8.

(2) Sinha, N.; Smith-Gill, J. S. Electrostatics in Protein Binding and Function. Current Protein & Peptide Science 2002, 3 (6), 601–614. https://doi.org/10.2174/1389203023380431.

(3) Nakamura, H. Roles of Electrostatic Interaction in Proteins. Quarterly Reviews of Biophysics 1996, 29 (1), 1–90. https://doi.org/10.1017/S0033583500005746.

(4) Zhou, H.-X.; Pang, X. Electrostatic Interactions in Protein Structure, Folding, Binding, and Condensation. Chem. Rev. 2018, 118 (4), 1691–1741. https://doi.org/10.1021/acs.chemrev.7b00305.

(5) Olsson, M. H. M.; Søndergaard, C. R.; Rostkowski, M.; Jensen, J. H. PROPKA3: Consistent Treatment of Internal and Surface Residues in Empirical PKa Predictions. J. Chem. Theory Comput. 2011, 7 (2), 525–537. https://doi.org/10.1021/ct100578z.

(6) Hofer, F.; Kraml, J.; Kahler, U.; Kamenik, A. S.; Liedl, K. R. Catalytic Site PKa Values of Aspartic, Cysteine, and Serine Proteases: Constant PH MD Simulations. J. Chem. Inf. Model. 2020. https://doi.org/10.1021/acs.jcim.0c00190.

(7) Gilson, M. K.; Given, J. A.; Bush, B. L.; McCammon, J. A. The Statistical-Thermodynamic Basis for Computation of Binding Affinities: A Critical Review. Biophys J 1997, 72 (3), 1047–1069. https://doi.org/10.1016/S0006-3495(97)78756-3.

(8) Mardis, K.; Luo, R.; David, L.; Potter, M.; Glemza, A.; Payne, G.; Gilson, M. K. Modeling Molecular Recognition: Theory and Application. Journal of Biomolecular Structure and Dynamics 2000, 17 (sup1), 89–94. https://doi.org/10.1080/07391102.2000.10506608.

(9) Kahler, U.; Kamenik, A. S.; Waibl, F.; Kraml, J.; Liedl, K. R. Protein-Protein Binding as a Two-Step Mechanism: Preselection of Encounter Poses during the Binding of BPTI and Trypsin. Biophysical Journal 2020, 119 (3), 652–666. https://doi.org/10.1016/j.bpj.2020.06.032.

(10) Waldner, B. J.; Kraml, J.; Kahler, U.; Spinn, A.; Schauperl, M.; Podewitz, M.; Fuchs, J. E.; Cruciani, G.; Liedl, K. R. Electrostatic Recognition in Substrate Binding to Serine Proteases. Journal of Molecular Recognition 2018, 31 (10), e2727. https://doi.org/10.1002/jmr.2727.

(11) Sinha, N.; Mohan, S.; Lipschultz, C. A.; Smith-Gill, S. J. Differences in Electrostatic Properties at Antibody–Antigen Binding Sites: Implications for Specificity and Cross-Reactivity. Biophysical Journal 2002, 83 (6), 2946–2968. https://doi.org/10.1016/S0006-3495(02)75302-2.

(12) Hofer, F.; Kamenik, A. S.; Fernández-Quintero, M. L.; Kraml, J.; Liedl, K. R. PH-Induced Local Unfolding of the Phl p 6 Pollen Allergen From CpH-MD. Frontiers in Molecular Biosciences 2021, 7, 477. https://doi.org/10.3389/fmolb.2020.603644.

(13) Chiu, M. L.; Goulet, D. R.; Teplyakov, A.; Gilliland, G. L. Antibody Structure and Function: The Basis for Engineering Therapeutics. Antibodies (Basel) 2019, 8 (4), 55. https://doi.org/10.3390/antib8040055.

(14) Kaplon, H.; Chenoweth, A.; Crescioli, S.; Reichert, J. M. Antibodies to Watch in 2022. Null 2022, 14 (1), 2014296. https://doi.org/10.1080/19420862.2021.2014296.

(15) Waibl, F.; Fernández-Quintero, M. L.; Kamenik, A. S.; Kraml, J.; Hofer, F.; Kettenberger, H.; Georges, G.; Liedl, K. R. Conformational Ensembles of Antibodies Determine Their Hydrophobicity. Biophysical Journal 2020. https://doi.org/10.1016/j.bpj.2020.11.010.

(16) Waibl, F.; Fernández-Quintero, M. L.; Wedl, F. S.; Kettenberger, H.; Georges, G.; Liedl, K. R. Comparison of Hydrophobicity Scales for Predicting Biophysical Properties of Antibodies. Frontiers in Molecular Biosciences 2022, 9.

(17) Kraft, T. E.; Richter, W. F.; Emrich, T.; Knaupp, A.; Schuster, M.; Wolfert, A.; Kettenberger, H. Heparin Chromatography as an in Vitro Predictor for Antibody Clearance Rate through Pinocytosis. mAbs 2020, 12 (1), 1683432. https://doi.org/10.1080/19420862.2019.1683432.

(18) Lienqueo, M. E.; Mahn, A.; Navarro, G.; Salgado, J. C.; Perez-Acle, T.; Rapaport, I.; Asenjo, J. A. New Approaches for Predicting Protein Retention Time in Hydrophobic Interaction Chromatography. Journal of Molecular Recognition 2006, 19 (4), 260–269. https://doi.org/10.1002/jmr.776.

(19) Holst, M.; Saied, F. Multigrid Solution of the Poisson—Boltzmann Equation. Journal of Computational Chemistry 1993, 14 (1), 105–113. https://doi.org/10.1002/jcc.540140114.

(20) Mongan, J.; Simmerling, C.; McCammon, J. A.; Case, D. A.; Onufriev, A. Generalized Born Model with a Simple, Robust Molecular Volume Correction. J. Chem. Theory Comput. 2007, 3 (1), 156–169. https://doi.org/10.1021/ct600085e.

(21) Izadi, S.; Harris, R. C.; Fenley, M. O.; Onufriev, A. V. Accuracy Comparison of Generalized Born Models in the Calculation of Electrostatic Binding Free Energies. J. Chem. Theory Comput. 2018, 14 (3), 1656–1670. https://doi.org/10.1021/acs.jctc.7b00886.

(22) Schrodinger. The AxPyMOL Molecular Graphics Plugin for Microsoft PowerPoint, Version 1.8, 2015.

(23) Humphrey, W.; Dalke, A.; Schulten, K. VMD: Visual Molecular Dynamics. Journal of Molecular Graphics 1996, 14 (1), 33–38. https://doi.org/10.1016/0263-7855(96)00018-5.

(24) Lijnzaad, P.; Berendsen, H. J. C.; Argos, P. Hydrophobic Patches on the Surfaces of Protein Structures. Proteins: Structure, Function, and Bioinformatics 1996, 25 (3), 389–397. https://doi.org/10.1002/(SICI)1097-0134(199607)25:3<389::AID-PROT10>3.0.CO;2-E.

(25) Lijnzaad, P.; Berendsen, H. J. C.; Argos, P. A Method for Detecting Hydrophobic Patches on Protein Surfaces. Proteins: Structure, Function, and Bioinformatics 1996, 26 (2), 192–203. https://doi.org/10.1002/(SICI)1097-0134(199610)26:2<192::AID-PROT9>3.0.CO;2-I.

(26) Molecular Operating Environment (MOE), 2020.

(27) Jain, T.; Sun, T.; Durand, S.; Hall, A.; Houston, N. R.; Nett, J. H.; Sharkey, B.; Bobrowicz, B.; Caffry, I.; Yu, Y.; Cao, Y.; Lynaugh, H.; Brown, M.; Baruah, H.; Gray, L. T.; Krauland, E. M.; Xu, Y.; Vásquez, M.; Wittrup, K. D. Biophysical Properties of the Clinical-Stage Antibody Landscape. Proc Natl Acad Sci USA 2017, 114 (5), 944. https://doi.org/10.1073/pnas.1616408114.

(28) Ruffolo, J. A.; Chu, L.-S.; Mahajan, S. P.; Gray, J. J. Fast, Accurate Antibody Structure Prediction from Deep Learning on Massive Set of Natural Antibodies. bioRxiv 2022, 2022.04.20.488972. https://doi.org/10.1101/2022.04.20.488972.

(29) Abanades, B.; Wong, W. K.; Boyles, F.; Georges, G.; Bujotzek, A.; Deane, C. M. ImmuneBuilder: Deep-Learning Models for Predicting the Structures of Immune Proteins. bioRxiv 2022, 2022.11.04.514231. https://doi.org/10.1101/2022.11.04.514231.

(30) Jurrus, E.; Engel, D.; Star, K.; Monson, K.; Brandi, J.; Felberg, L. E.; Brookes, D. H.; Wilson, L.; Chen, J.; Liles, K.; Chun, M.; Li, P.; Gohara, D. W.; Dolinsky, T.; Konecny, R.; Koes, D. R.; Nielsen, J. E.; Head-Gordon, T.; Geng, W.; Krasny, R.; Wei, G.-W.; Holst, M. J.; McCammon, J. A.; Baker, N. A. Improvements to the APBS Biomolecular Solvation Software Suite. Protein Science 2018, 27 (1), 112–128. https://doi.org/10.1002/pro.3280.

(31) Baker, N. A.; Sept, D.; Joseph, S.; Holst, M. J.; McCammon, J. A. Electrostatics of Nanosystems: Application to Microtubules and the Ribosome. Proceedings of the National Academy of Sciences 2001, 98 (18), 10037–10041. https://doi.org/10.1073/pnas.181342398.

(32) Schrodinger. The PyMOL Molecular Graphics System, Version 1.8, 2015.

(33) López-Otín, C.; Bond, J. S. Proteases: Multifunctional Enzymes in Life and Disease. J Biol Chem 2008, 283 (45), 30433–30437. https://doi.org/10.1074/jbc.R800035200.

(34) Puente, X. S.; Sánchez, L. M.; Gutiérrez-Fernández, A.; Velasco, G.; López-Otín, C. A Genomic View of the Complexity of Mammalian Proteolytic Systems. Biochemical Society Transactions 2005, 33 (2), 331–334. https://doi.org/10.1042/BST0330331.

(35) Fuchs, J. E.; Liedl, K. R. Substrate Sequences Tell Similar Stories as Binding Cavities: Commentary. J. Chem. Inf. Model. 2013, 53 (12), 3115–3116. https://doi.org/10.1021/ci4005783.

(36) Fuchs, J. E.; Huber, R. G.; Waldner, B. J.; Kahler, U.; von Grafenstein, S.; Kramer, C.; Liedl, K. R. Dynamics Govern Specificity of a Protein-Protein Interface: Substrate Recognition by Thrombin. PLOS ONE 2015, 10 (10), e0140713. https://doi.org/10.1371/journal.pone.0140713.

(37) Rawlings, N. D.; Barrett, A. J.; Finn, R. Twenty Years of the MEROPS Database of Proteolytic Enzymes, Their Substrates and Inhibitors. Nucleic Acids Research 2016, 44 (D1), D343–D350. https://doi.org/10.1093/nar/gkv1118.

(38) Tu, C.; Terraube, V.; Tam, A. S. P.; Stochaj, W.; Fennell, B. J.; Lin, L.; Stahl, M.; LaVallie, E. R.; Somers, W.; Finlay, W. J. J.; Mosyak, L.; Bard, J.; Cunningham, O. A Combination of Structural and Empirical Analyses Delineates the Key Contacts Mediating Stability and Affinity Increases in an Optimized Biotherapeutic Single-Chain Fv (ScFv)*. Journal of Biological Chemistry 2016, 291 (3), 1267–1276. https://doi.org/10.1074/jbc.M115.688010.

(39) Foote, J.; Winter, G. Antibody Framework Residues Affecting the Conformation of the Hypervariable Loops. Journal of Molecular Biology 1992, 224 (2), 487–499. https://doi.org/10.1016/0022-2836(92)91010-M.

(40) Tramontano, A.; Chothia, C.; Lesk, A. M. Framework Residue 71 Is a Major Determinant of the Position and Conformation of the Second Hypervariable Region in the VH Domains of Immunoglobulins. Journal of Molecular Biology 1990, 215 (1), 175–182. https://doi.org/10.1016/S0022-2836(05)80102-0.

(41) Dunbar, J.; Fuchs, A.; Shi, J.; Deane, C. M. ABangle: Characterising the VH–VL Orientation in Antibodies. Protein Engineering, Design and Selection 2013, 26 (10), 611–620. https://doi.org/10.1093/protein/gzt020.

(42) Fernández-Quintero, M. L.; Hoerschinger, V. J.; Lamp, L. M.; Bujotzek, A.; Georges, G.; Liedl, K. R. VH-VL Interdomain Dynamics Observed by Computer Simulations and NMR. Proteins: Structure, Function, and Bioinformatics 2020, n/a (n/a). https://doi.org/10.1002/prot.25872.

(43) Fernández-Quintero, M. L.; Kroell, K. B.; Hofer, F.; Riccabona, J. R.; Liedl, K. R. Mutation of Framework Residue H71 Results in Different Antibody Paratope States in Solution. Frontiers in Immunology 2021, 12, 243. https://doi.org/10.3389/fimmu.2021.630034.

(44) Krauss, J.; Arndt, M. A. E.; Zhu, Z.; Newton, D. L.; Vu, B. K.; Choudhry, V.; Darbha, R.; Ji, X.; Courtenay-Luck, N. S.; Deonarain, M. P.; Richards, J.; Rybak, S. M. Impact of Antibody Framework Residue VH-71 on the Stability of a Humanised Anti-MUC1 ScFv and Derived Immunoenzyme. British Journal of Cancer 2004, 90 (9), 1863–1870. https://doi.org/10.1038/sj.bjc.6601759.

(45) Honegger, A.; Malebranche, A. D.; Röthlisberger, D.; Plückthun, A. The Influence of the Framework Core Residues on the Biophysical Properties of Immunoglobulin Heavy Chain Variable Domains. Protein Engineering, Design and Selection 2009, 22 (3), 121–134. https://doi.org/10.1093/protein/gzn077.

(46) Dunbar, J.; Deane, C. M. ANARCI: Antigen Receptor Numbering and Receptor Classification. Bioinformatics (Oxford, England) 2016, 32 (2), 298–300. https://doi.org/10.1093/bioinformatics/btv552.

(47) Nguyen, C. N.; Young, T. K.; Gilson, M. K. Grid Inhomogeneous Solvation Theory: Hydration Structure and Thermodynamics of the Miniature Receptor Cucurbit[7]Uril. J Chem Phys 2012, 137 (4), 044101–044101. https://doi.org/10.1063/1.4733951.

(48) Kraml, J.; Kamenik, A. S.; Waibl, F.; Schauperl, M.; Liedl, K. R. Solvation Free Energy as a Measure of Hydrophobicity: Application to Serine Protease Binding Interfaces. J. Chem. Theory Comput. 2019, 15 (11), 5872–5882. https://doi.org/10.1021/acs.jctc.9b00742.

(49) Schauperl, M.; Podewitz, M.; Waldner, B. J.; Liedl, K. R. Enthalpic and Entropic Contributions to Hydrophobicity. J Chem Theory Comput 2016, 12 (9), 4600–4610. https://doi.org/10.1021/acs.jctc.6b00422.

(50) Waibl, F.; Fernández-Quintero, M. L.; Kamenik, A. S.; Kraml, J.; Hofer, F.; Kettenberger, H.; Georges, G.; Liedl, K. R. Conformational Ensembles of Antibodies Determine Their Hydrophobicity. Biophysical Journal 2021, 120 (1), 143–157. https://doi.org/10.1016/j.bpj.2020.11.010.

