## Supporting Information for "PEP-Patch: Electrostatics in Protein-Protein Recognition, Specificity and Antibody Developability"

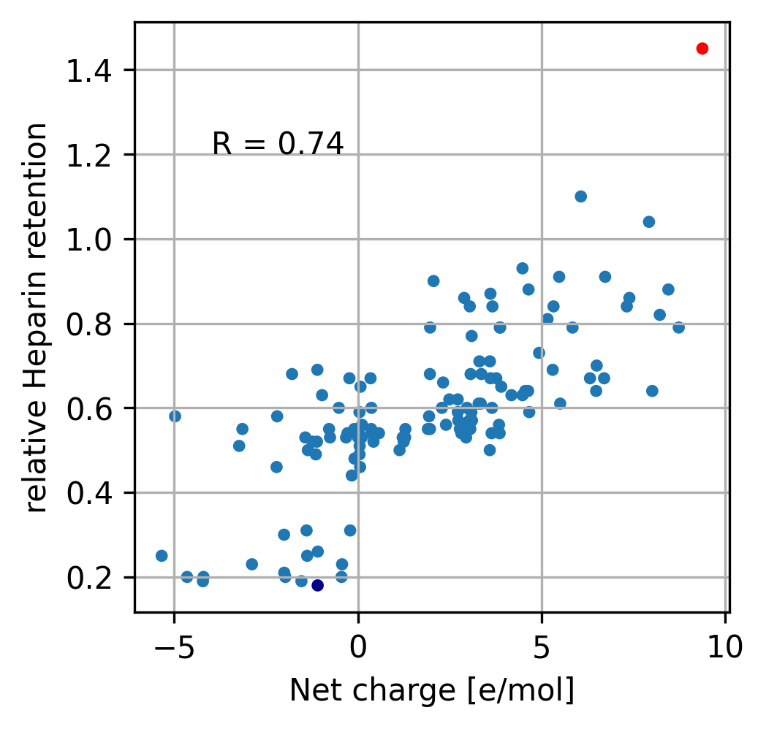

Figure S1: Relative heparin retention times correlated with the positive patch area provided by MOE. As in Figure 3 of the main text, lenzilumab and sirukumab are shown as red and blue dots, respectively.

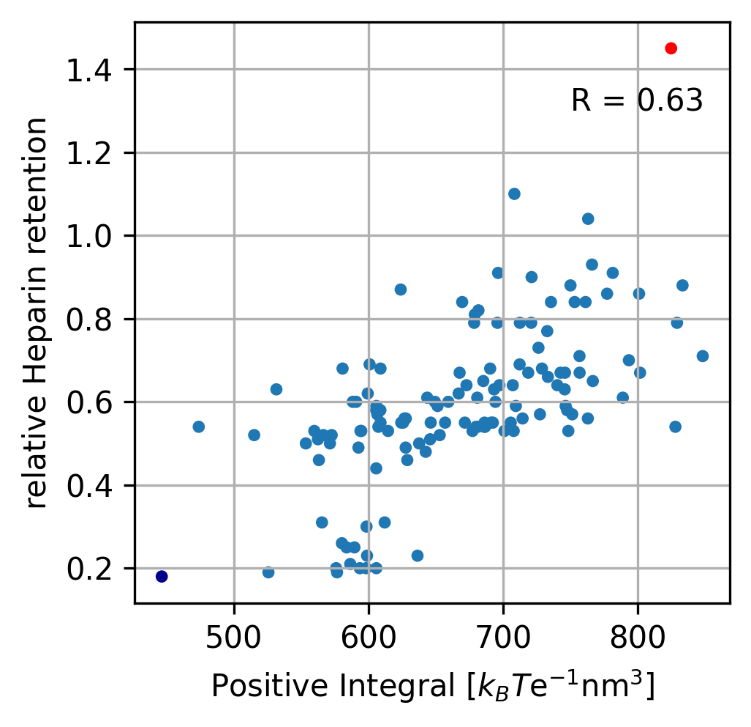

Figure S2: Net charge of the antibodies calculated with MOE, correlated with the relative heparin retention time of the antibodies.

Table S1: Residues contributing the most to the absolute value ascribed to an electrostatic patch for the results presented in Figure 1 and Figure 2.

| Trypsin | | | |
| --- | --- | --- | --- |
| type | npoints | area | main_residue |
| positive | 729 | 117.679463 | LYS188 |
| positive | 721 | 116.802802 | ARG166 |
| positive | 404 | 65.1777161 | ARG88 |
| positive | 339 | 55.5023517 | LEU73 |
| positive | 314 | 51.9860782 | ARG119 |
| positive | 312 | 51.1179195 | PRO91 |
| positive | 274 | 42.6745675 | ARG35 |
| positive | 233 | 36.3607999 | ARG75 |
| positive | 157 | 27.3872189 | SER83 |
| positive | 156 | 24.2751487 | ARG69 |
| positive | 121 | 20.8056815 | THR76 |
| positive | 69 | 10.5907007 | SER147 |
| positive | 64 | 10.2707108 | VAL89 |
| positive | 45 | 6.38613449 | GLY37 |
| positive | 38 | 6.00178771 | GLN63 |
| positive | 37 | 5.95083239 | SER123 |
| positive | 38 | 5.21988658 | TRP138 |
| positive | 15 | 2.52468883 | THR164 |
| positive | 13 | 2.48472098 | SER184 |
| positive | 16 | 2.23689742 | LEU231 |
| positive | 13 | 2.05111792 | ASN151 |
| positive | 11 | 1.64536576 | THR171 |
| positive | 14 | 1.2597704 | ARG69 |
| positive | 6 | 0.61085037 | SER172 |
| positive | 5 | 0.26974551 | GLY19 |
| positive | 6 | 0.15791531 | GLY37 |
| positive | 6 | 0.12617516 | GLY37 |
| positive | 10 | 0.09310352 | SER72 |
| positive | 6 | 0.08882306 | LYS154 |
| positive | 6 | 0.07800999 | SER32 |
| positive | 6 | 0.06634554 | ARG119 |
| positive | 6 | 0.01333066 | THR156 |
| positive | 6 | 0.00865386 | TRP138 |
| positive | 6 | 0.00497744 | LYS188 |
| negative | 860 | 138.922415 | GLN192 |
| negative | 274 | 45.585308 | ASP26 |
| negative | 109 | 17.5881279 | ALA239 |
| negative | 84 | 11.7466516 | GLY182 |
| negative | 64 | 8.16579672 | ASP201 |
| negative | 29 | 5.18963711 | SER59 |
| negative | 31 | 4.5760538 | SER222 |
| negative | 22 | 1.99737441 | TYR221 |
| negative | 8 | 1.22504268 | PHE179 |
| negative | 7 | 1.17262072 | ASP26 |
| negative | 10 | 0.92894108 | ALA239 |
| negative | 5 | 0.89138293 | ALA239 |
| negative | 5 | 0.71569338 | ALA173 |
| negative | 14 | 0.60340463 | VAL224 |
| negative | 6 | 0.1454201 | ALA209 |
| negative | 6 | 0.13372595 | MET178 |
| negative | 6 | 0.09828762 | TYR221 |
| negative | 6 | 0.09431558 | ALA209 |
| negative | 6 | 0.06020165 | ALA209 |
| negative | 6 | 0.03681595 | SER190 |

| Chymotrypsin | | | |
| --- | --- | --- | --- |
| type | npoints | area | main_residue |
| positive | 7943 | 1312.55305 | LYS93 |
| positive | 793 | 129.316283 | CYS1 |
| positive | 657 | 106.886058 | LYS84 |
| positive | 567 | 89.8772885 | LYS202 |
| positive | 224 | 33.8326658 | SER190 |
| positive | 84 | 11.061201 | SER189 |
| positive | 61 | 9.4477061 | ARG145 |
| positive | 49 | 8.27085818 | LYS36 |
| positive | 38 | 6.65437407 | THR222 |
| positive | 42 | 5.22824885 | THR138 |
| positive | 27 | 4.13088373 | SER119 |
| positive | 30 | 3.43381332 | LEU123 |
| positive | 17 | 2.82974537 | LYS203 |
| positive | 21 | 2.76323791 | ALA126 |
| positive | 14 | 2.25412412 | LYS82 |
| positive | 16 | 2.0558185 | GLY140 |
| positive | 15 | 1.86168774 | VAL210 |
| positive | 7 | 1.53223932 | LYS203 |
| positive | 18 | 1.31616817 | VAL238 |
| positive | 2 | 0.40298813 | SER45 |
| positive | 4 | 0.35489217 | ALA56 |
| positive | 2 | 0.31336404 | SER218 |
| positive | 6 | 0.12671322 | THR232 |
| positive | 6 | 0.12182391 | VAL231 |
| positive | 1 | 0.11814837 | LYS90 |
| positive | 6 | 0.1179208 | SER190 |
| positive | 6 | 0.06820048 | VAL60 |
| positive | 6 | 0.02323131 | VAL3 |
| positive | 6 | 0.02227583 | GLY226 |
| positive | 6 | 0.01481454 | TRP237 |
| positive | 6 | 0.00013212 | GLY226 |
| negative | 7772 | 1284.84966 | PHE39 |
| negative | 217 | 36.6977191 | TYR146 |
| negative | 112 | 18.6612543 | ASN245 |
| negative | 107 | 18.3547825 | GLU49 |
| negative | 21 | 3.03103765 | ASP194 |
| negative | 18 | 2.72215966 | ASP35 |
| negative | 19 | 2.53942174 | GLY196 |
| negative | 13 | 1.2697544 | SER214 |
| negative | 10 | 0.74375484 | TRP29 |
| negative | 2 | 0.54043506 | SER127 |
| negative | 6 | 0.37061736 | ILE85 |
| negative | 4 | 0.27287853 | ASP128 |
| negative | 6 | 0.27247725 | VAL67 |
| negative | 6 | 0.26031518 | GLY25 |
| negative | 1 | 0.16794804 | ASP35 |
| negative | 6 | 0.14523811 | THR139 |
| negative | 6 | 0.12813773 | GLY25 |
| negative | 6 | 0.09596294 | SER214 |
| negative | 10 | 0.06544949 | THR138 |
| negative | 6 | 0.02181548 | VAL23 |
| negative | 6 | 0.00540477 | ASP102 |
| negative | 6 | 9.52E-05 | LEU33 |

| Granzyme B | | | |
| --- | --- | --- | --- |
| type | npoints | area | main_residue |
| positive | 13528 | 2537.12056 | LYS188 |
| positive | 6210 | 1166.51848 | LYS192 |
| positive | 1715 | 319.434124 | ARG110 |
| positive | 284 | 52.747294 | LYS131 |
| positive | 266 | 49.4404739 | ARG27 |
| positive | 256 | 47.4218536 | GLN210 |
| positive | 172 | 34.12302 | ARG87 |
| positive | 181 | 34.0747797 | ARG217 |
| positive | 62 | 12.8301405 | GLN37 |
| positive | 50 | 9.00056703 | HIS153 |
| positive | 48 | 8.33189323 | ALA139 |
| positive | 40 | 7.49171164 | LYS149 |
| positive | 42 | 5.35365681 | TRP141 |
| positive | 25 | 4.17261904 | THR144 |
| positive | 8 | 1.68319615 | ASN219 |
| positive | 12 | 1.58704484 | GLN129 |
| positive | 6 | 1.50931899 | ARG172 |
| positive | 15 | 1.32892487 | VAL52 |
| positive | 14 | 1.18532356 | ARG41 |
| positive | 16 | 1.09542089 | GLY193 |
| positive | 4 | 0.89515589 | ASN219 |
| positive | 6 | 0.66125942 | GLN156 |
| positive | 6 | 0.39810759 | MET224 |
| positive | 6 | 0.39638472 | TYR32 |
| positive | 5 | 0.3301109 | GLN163 |
| positive | 2 | 0.32290892 | TYR245 |
| positive | 6 | 0.25931213 | ARG41 |
| positive | 6 | 0.20154489 | THR189 |
| positive | 1 | 0.1994544 | ARG87 |
| positive | 3 | 0.14950518 | PRO120 |
| positive | 6 | 0.10986478 | THR144 |
| positive | 5 | 0.10781241 | PRO120 |
| positive | 6 | 0.10062683 | ALA31 |
| positive | 6 | 0.0899475 | VAL52 |
| positive | 6 | 0.06556522 | ARG41 |
| positive | 6 | 0.04740378 | GLN210 |
| positive | 6 | 0.04277298 | VAL158 |
| positive | 1 | 0.03983786 | PHE82 |
| positive | 6 | 0.02321586 | GLY43 |
| positive | 6 | 0.01928432 | ASP49 |
| positive | 6 | 0.01655484 | VAL130 |
| positive | 6 | 0.01208086 | LEU33 |
| positive | 1 | 0.00983955 | PRO146 |
| positive | 6 | 0.00869647 | LEU171 |
| positive | 6 | 0.00633369 | VAL235 |
| positive | 6 | 0.00511543 | LYS188 |
| positive | 1 | 0.00385118 | ARG226 |
| positive | 6 | 5.90E-05 | LYS239 |
| negative | 1941 | 365.909166 | ASP176 |
| negative | 773 | 143.496239 | GLU75 |
| negative | 191 | 35.0881447 | TYR245 |
| negative | 108 | 19.0411565 | GLU186 |
| negative | 56 | 8.16209073 | LEU181 |
| negative | 35 | 5.87209033 | ASP37 |
| negative | 27 | 4.24344888 | TYR245 |
| negative | 19 | 3.86103188 | ASP50 |
| negative | 16 | 3.83266122 | GLU109 |
| negative | 10 | 1.3100794 | PHE51 |
| negative | 14 | 1.16955116 | VAL130 |
| negative | 4 | 0.33034444 | VAL52 |
| negative | 6 | 0.32864457 | ALA112 |
| negative | 1 | 0.25826574 | TRP59 |
| negative | 5 | 0.24752879 | SER190 |
| negative | 1 | 0.16138442 | GLU109 |
| negative | 5 | 0.11500604 | ARG114 |
| negative | 1 | 0.07741278 | ASP49 |
| negative | 1 | 0.06624385 | CYS228 |
| negative | 3 | 0.06390481 | ARG114 |
| negative | 6 | 0.0516718 | MET242 |
| negative | 6 | 0.03608713 | PRO28 |
| negative | 6 | 0.03530829 | VAL130 |
| negative | 6 | 0.03212897 | TYR245 |
| negative | 6 | 0.02849412 | GLN129 |
| negative | 6 | 0.0101261 | SER100 |
| negative | 6 | 0.00997878 | VAL162 |
| negative | 6 | 0.00726417 | GLU75 |
| negative | 6 | 0.0065888 | ALA112 |
| negative | 6 | 0.00256342 | PRO225 |

| 3b4 | | | |
| --- | --- | --- | --- |
| type | npoints | area | main_residue |
| positive | 2512 | 428.789227 | LYS62 |
| positive | 924 | 157.953631 | LYS13 |
| positive | 716 | 122.127738 | LYS45 |
| positive | 538 | 91.4414097 | GLN1 |
| positive | 364 | 60.4605843 | ARG94 |
| positive | 101 | 16.3860577 | LYS23 |
| positive | 95 | 15.4982389 | LYS66 |
| positive | 55 | 7.83732517 | THR68 |
| positive | 44 | 7.15705111 | LYS42 |
| positive | 46 | 6.5778703 | ASN52 |
| positive | 29 | 5.35951325 | ASN69 |
| positive | 38 | 5.05948052 | LYS62 |
| positive | 30 | 4.8667855 | SER67 |
| positive | 24 | 4.43226678 | SER27 |
| positive | 22 | 4.41414826 | HIS60 |
| positive | 20 | 3.06133387 | ARG94 |
| positive | 26 | 2.66122019 | VAL51 |
| positive | 11 | 1.80785497 | ARG94 |
| positive | 5 | 1.3310678 | ARG54 |
| positive | 9 | 1.31000165 | THR108 |
| positive | 8 | 1.19027251 | SER27 |
| positive | 5 | 0.82715504 | ARG93 |
| positive | 3 | 0.53213758 | SER84 |
| positive | 6 | 0.29187666 | GLN1 |
| positive | 1 | 0.27476218 | GLY57 |
| positive | 1 | 0.25447908 | SER82 |
| positive | 2 | 0.2473777 | LYS103 |
| positive | 5 | 0.24251281 | SER27 |
| positive | 1 | 0.16593153 | LYS12 |
| positive | 6 | 0.14984186 | PHE63 |
| positive | 1 | 0.12056577 | ALA43 |
| positive | 1 | 0.07442074 | PRO52 |
| positive | 6 | 0.05451902 | THR68 |
| positive | 6 | 0.023932 | ALA71 |
| negative | 6259 | 1064.54774 | TYR100 |
| negative | 2285 | 393.890378 | GLU108 |
| negative | 1007 | 170.014341 | ASP85 |
| negative | 662 | 114.060594 | GLU73 |
| negative | 547 | 93.2590157 | SER113 |
| negative | 150 | 26.3165134 | PRO96 |
| negative | 94 | 17.3531439 | ASP72 |
| negative | 42 | 6.88287011 | GLN1 |
| negative | 40 | 6.84576262 | THR74 |
| negative | 29 | 4.92600511 | ASP27 |
| negative | 28 | 4.52016513 | PRO15 |
| negative | 19 | 3.67606112 | LYS62 |
| negative | 17 | 2.43431612 | GLU10 |
| negative | 6 | 0.90466735 | GLU10 |
| negative | 7 | 0.61822932 | SER30 |
| negative | 10 | 0.48974506 | ASP50 |
| negative | 10 | 0.16774348 | ASP50 |
| negative | 6 | 0.10199716 | TRP32 |
| negative | 6 | 0.0664382 | TYR98 |
| negative | 6 | 0.0490952 | ASP100 |
| negative | 6 | 0.04611165 | SER89 |
| negative | 6 | 0.0220922 | TYR91 |
| negative | 6 | 0.01229754 | PHE49 |
| negative | 6 | 0.00750903 | SER34 |
| negative | 6 | 0.00624069 | ASN58 |
| negative | 6 | 5.45E-07 | LEU78 |

| e10 | | | |
| --- | --- | --- | --- |
| type | npoints | area | main_residue |
| positive | 1268 | 203.225349 | SER30 |
| positive | 946 | 150.861561 | LYS62 |
| positive | 613 | 101.975944 | GLN1 |
| positive | 611 | 97.6618333 | LYS13 |
| positive | 594 | 94.4855458 | HIS39 |
| positive | 375 | 59.9213342 | ARG83 |
| positive | 346 | 55.8554497 | LYS23 |
| positive | 184 | 30.1797673 | SER84 |
| positive | 96 | 15.2559337 | ARG61 |
| positive | 87 | 13.6223383 | ARG54 |
| positive | 90 | 13.2826421 | TRP103 |
| positive | 53 | 8.70396899 | ARG54 |
| positive | 47 | 7.84472203 | SER18 |
| positive | 49 | 7.6503087 | LYS23 |
| positive | 56 | 6.7328299 | SER84 |
| positive | 44 | 6.44911319 | TRP47 |
| positive | 31 | 4.74468231 | SER27 |
| positive | 29 | 4.55253409 | ARG83 |
| positive | 17 | 2.51942562 | THR105 |
| positive | 13 | 2.37319543 | SER12 |
| positive | 14 | 2.15572966 | THR28 |
| positive | 23 | 2.08518759 | ALA13 |
| positive | 6 | 1.05484618 | ASN69 |
| positive | 6 | 0.77171049 | LEU94 |
| positive | 7 | 0.59753573 | TYR79 |
| positive | 6 | 0.46724068 | ARG66 |
| positive | 3 | 0.44333242 | SER18 |
| positive | 5 | 0.42644175 | ILE51 |
| positive | 5 | 0.37793912 | LYS19 |
| positive | 6 | 0.30603493 | PHE63 |
| positive | 1 | 0.2296781 | ASN69 |
| positive | 1 | 0.22690341 | SER82 |
| positive | 1 | 0.22336508 | LYS62 |
| positive | 6 | 0.19701217 | GLY49 |
| positive | 10 | 0.1763086 | GLY49 |
| positive | 1 | 0.1558938 | SER25 |
| positive | 1 | 0.13864917 | SER2 |
| positive | 6 | 0.06627006 | TYR32 |
| positive | 6 | 0.06355005 | LEU4 |
| positive | 6 | 0.0617888 | LYS62 |
| positive | 6 | 0.05808696 | GLN1 |
| positive | 1 | 0.05680838 | SER82 |
| positive | 6 | 0.05569998 | ILE51 |
| positive | 1 | 0.03626656 | SER27 |
| positive | 6 | 0.03074973 | ILE69 |
| positive | 6 | 0.02795613 | TYR36 |
| positive | 6 | 0.01597983 | MET48 |
| positive | 6 | 3.34E-05 | VAL37 |
| negative | 7521 | 1214.99069 | TYR100 |
| negative | 3273 | 526.63814 | LEU106 |
| negative | 760 | 122.388021 | GLU73 |
| negative | 408 | 68.0369264 | GLU46 |
| negative | 407 | 61.8737191 | GLN43 |
| negative | 279 | 45.2769291 | THR68 |
| negative | 206 | 33.1061826 | SER113 |
| negative | 155 | 24.5935023 | ASP82 |
| negative | 130 | 20.1110056 | GLU81 |
| negative | 107 | 17.180903 | GLU10 |
| negative | 96 | 15.0344981 | ASP72 |
| negative | 69 | 11.1367083 | SER2 |
| negative | 54 | 8.69032998 | GLN3 |
| negative | 50 | 7.81803451 | VAL51 |
| negative | 38 | 4.79547475 | SER25 |
| negative | 18 | 3.19399885 | GLN3 |
| negative | 21 | 3.16127543 | LEU45 |
| negative | 13 | 2.33366788 | ASP27 |
| negative | 7 | 0.85954202 | GLN43 |
| negative | 6 | 0.45500448 | LEU47 |
| negative | 5 | 0.33013279 | SER14 |
| negative | 1 | 0.22811838 | ASP100 |
| negative | 4 | 0.19204181 | GLY99 |
| negative | 5 | 0.17999254 | ALA43 |
| negative | 10 | 0.13938728 | ASP95 |
| negative | 1 | 0.1148977 | PRO55 |
| negative | 1 | 0.01698894 | GLN43 |
| negative | 6 | 0.01377142 | VAL58 |
| negative | 6 | 0.01110618 | VAL33 |
| negative | 6 | 0.0011787 | ILE75 |

Table S2: PDB accession codes of the 49 available crystal structures of the Jain et al. dataset. [PNAS 2017 Vol. 114 Issue 5 Pages 944-949 DOI: 10.1073/pnas.1616408114]

| **Antibody** | **PDB code** |
| --- | --- |
| abituzumab |  |
| abrilumab |  |
| adalimumab | 4nyl |
| alemtuzumab | 1bey |
| anifrolumab | 4qxg |
| atezolizumab | 5x8l |
| bapineuzumab | 4ojf |
| basiliximab | 1mim |
| belimumab | 5y9j |
| benralizumab |  |
| bevacizumab | 1bj1 |
| bimagrumab | 5ngv |
| blosozumab |  |
| bococizumab |  |
| briakinumab | 5n2k |
| brodalumab |  |
| canakinumab | 4g6j |
| carlumab |  |
| certolizumab | 5wux |
| cetuximab | 1yy8 |
| cixutumumab |  |
| clazakizumab |  |
| codrituzumab |  |
| crenezumab | 5vzy |
| dacetuzumab |  |
| daclizumab | 3nfp |
| dalotuzumab |  |
| daratumumab |  |
| denosumab |  |
| drozitumab | 4od2 |
| duligotuzumab |  |
| dupilumab |  |
| eculizumab | 5i5k |
| efalizumab | 3eoa |
| eldelumab |  |
| elotuzumab |  |
| emibetuzumab |  |
| enokizumab |  |
| epratuzumab | 5vl3 |
| etrolizumab |  |
| evolocumab |  |
| fasinumab |  |
| fezakinumab |  |
| ficlatuzumab |  |
| fletikumab |  |
| foralumab |  |
| fresolimumab |  |
| fulranumab |  |
| galiximab |  |
| ganitumab |  |
| gantenerumab |  |
| gemtuzumab |  |
| gevokizumab | 4g6m |
| glembatumumab |  |
| golimumab | 5yoy |
| guselkumab |  |
| ibalizumab | 3o2d |
| imgatuzumab |  |
| infliximab | 4g3y |
| inotuzumab |  |
| ipilimumab | 5tru |
| ixekizumab | 6-Nov |
| lampalizumab |  |
| lebrikizumab | 4i77 |
| lenzilumab |  |
| lintuzumab |  |
| lirilumab |  |
| matuzumab | 3c09 |
| mavrilimumab |  |
| mepolizumab |  |
| mogamulizumab |  |
| motavizumab | 3ixt |
| muromonab | 1sy6 |
| natalizumab | 4irz |
| necitumumab | 6b3s |
| nimotuzumab |  |
| nivolumab | 5wt9 |
| obinutuzumab |  |
| ocrelizumab |  |
| ofatumumab | 3giz |
| olokizumab | 4cni |
| omalizumab | 2xa8 |
| onartuzumab | 4k3j |
| otelixizumab |  |
| otlertuzumab |  |
| ozanezumab |  |
| palivizumab |  |
| panitumumab | 5sx5 |
| panobacumab |  |
| parsatuzumab |  |
| patritumab |  |
| pembrolizumab | 5ggs |
| pertuzumab | 1s78 |
| pinatuzumab |  |
| polatuzumab |  |
| ponezumab | 3u0t |
| ramucirumab |  |
| ranibizumab | 1cz8 |
| reslizumab |  |
| rilotumumab |  |
| rituximab | 6vja |
| robatumumab |  |
| romosozumab |  |
| sarilumab |  |
| secukinumab |  |
| seribantumab |  |
| sifalimumab | 4ypg |
| siltuximab |  |
| simtuzumab |  |
| sirukumab |  |
| tabalumab |  |
| tanezumab | 4edw |
| teplizumab |  |
| tigatuzumab |  |
| tildrakizumab |  |
| tocilizumab |  |
| tovetumab |  |
| tralokinumab | 5l6y |
| trastuzumab | 1n8z |
| tremelimumab | 5ggv |
| urelumab | 6mhr |
| ustekinumab | 3hmw |
| vedolizumab |  |
| visilizumab |  |
| zalutumumab |  |
| zanolimumab |  |
